# Massively parallel sample preparation for multiplexed single-cell proteomics using nPOP

**DOI:** 10.1101/2023.11.27.568927

**Authors:** Andrew Leduc, Luke Koury, Joshua Cantlon, Nikolai Slavov

## Abstract

Single-cell proteomics by mass spectrometry (MS) allows quantifying proteins with high specificity and sensitivity. To increase its throughput, we developed nPOP, a method for parallel preparation of thousands of single cells in nanoliter volume droplets deposited on glass slides. Here, we describe its protocol with emphasis on its flexibility to prepare samples for different multiplexed MS methods. An implementation with plexDIA demonstrates accurate quantification of about 3,000 - 3,700 proteins per human cell. The protocol is implemented on the CellenONE instrument and uses readily available consumables, which should facilitate broad adoption. nPOP can be applied to all samples that can be processed to a single-cell suspension. It takes 1 or 2 days to prepare over 3,000 single cells. We provide metrics and software for quality control that can support the robust scaling of nPOP to higher plex reagents for achieving reliable high-throughput single-cell protein analysis.

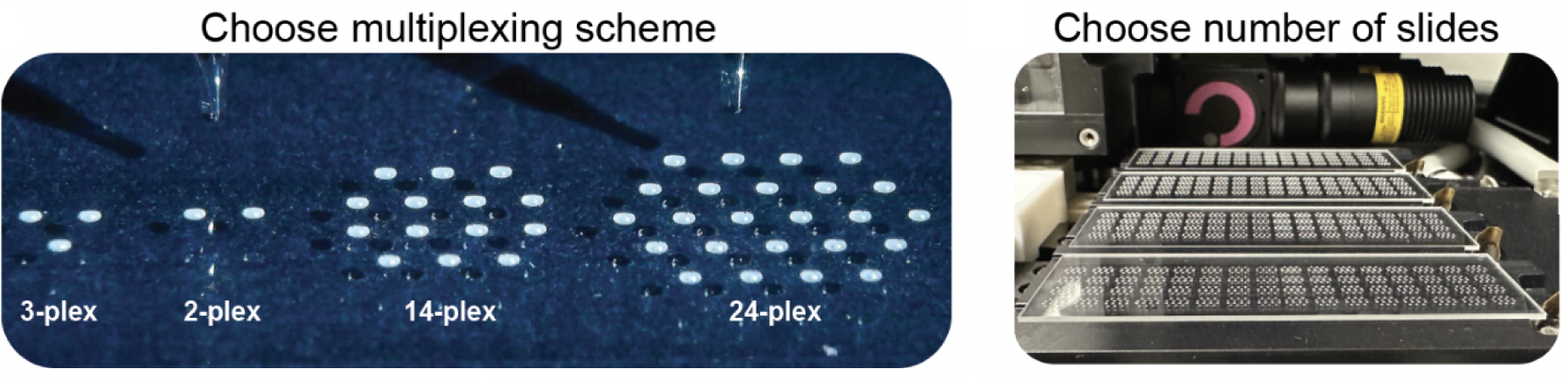

## Introduction

The application of single-cell technologies has advanced our understanding of how transcriptional regulation shapes and is shaped by the diverse cell types that comprise tissues. These advances have crucially depended on highly multiplexed molecular and droplet based library preparation methods for RNA sequencing. However, developing a more complete understanding of these cell types and their roles in organ function and dysfunction also requires methods for measuring protein abundance, modifications and interactions from many thousands of single cells with high specificity and accuracy^1,2^.

For decades, antibody based methods such as mass cytometry have enabled detecting epitopes from dozens of proteins at high throughput, but they are limited by the specificity of antibody binding, the depth of protein coverage, and the detectable epitopes^3^. More recently, mass spectrometry (MS) based proteomics has enabled specific and accurate quantification of thousands of proteins in single cells ^4–16^. These methods allow counting millions of peptide copies per single cell, which can support protein quantification with reliable count statistics^6,17,18^. Furthermore, single-cell MS proteomics can support the estimation of quantification reliability^7^ and functional protein analysis, such as protein conformations^19^, which can enable quantitative models and biological inferences^20^. However, currently the throughput of MS has been limited^21^. The limitation stems from the time needed to separate peptides, usually by liquid chromatography or capillary electrophoresis, and thus the time required for analyzing enough single cells.

Increases in the throughput of single-cell protein analysis by MS have taken two forms. First, with the introduction of methods for isolating and fragmenting multiple peptides in parallel ^22,23^, the amount of time required to identify thousands of proteins in a single MS run has significantly decreased from hours to 5 minutes ^24^. However, even at a throughput of one sample every five minutes, analyzing thousands of single cells would still take weeks. Second, multiplexing single cells utilizing isobaric or non-isobaric mass tags has enabled simultaneous measurement of proteins from multiple single cells in one MS run^17,25^. plexDIA combines the benefits of multiplexing and parallel peptide fragmentation, and 3-plexDIA of single cells on short 10 minute gradients has achieved throughput of 1 cell per 3⅓ minutes of active gradient^17^. The approach has the potential to significantly increase the scale of multiplexing^26^, which in turn requires methods for sample preparation that can easily prepare thousands of samples for multiplexed analysis. The method detailed in this protocol aims to meet this need and help further scale the throughput of single-cell proteomics^21^.

### Development of nPOP

#### Sample preparation

Leduc *et al.* developed nPOP to enable the preparation of thousands of high quality single-cell samples in an automated fashion for multiplexed analysis by LC/MS ^7^. We aimed to enable flexible experimental designs to accommodate different multiplexing schemes without requiring specialized consumables; nPOP supports all existing multiplexing approaches and can be easily programmed to new ones that may emerge. We also aimed to minimize reaction volumes^27^ while simplifying sample preparation and increasing its throughput.

nPOP achieves these goals by preparing single cells in droplets on the surface of unpatterned microscope glass slides, avoiding restrictions of predefined wells, Fig. 1. This design allows for minimizing reaction volumes of droplets to 10-20 nL and spatial freedom for droplet positioning based on the desired plex for multiplexing. The droplet positions are computationally programed, and the programmed position can be easily and quickly reprogramed. Together, these features enable increased spatial proximity of these droplets, maximizing the number of cells that can be prepared over the surface of the slide, Fig. 1a, and Fig. 2. Additionally, the small volumes allow single cell reactions to have a sufficient concentration of reactants while minimizing the total amount of reagent added, thus reducing reagent costs and waste. As a result, the total reagent cost is about $0.12 / cell, with detailed estimates provided in Table 1. This design is enabled by the spatial precision and picoliter dispensing capabilities of the CellenONE cell dispenser and liquid handler.

**Figure 1:**
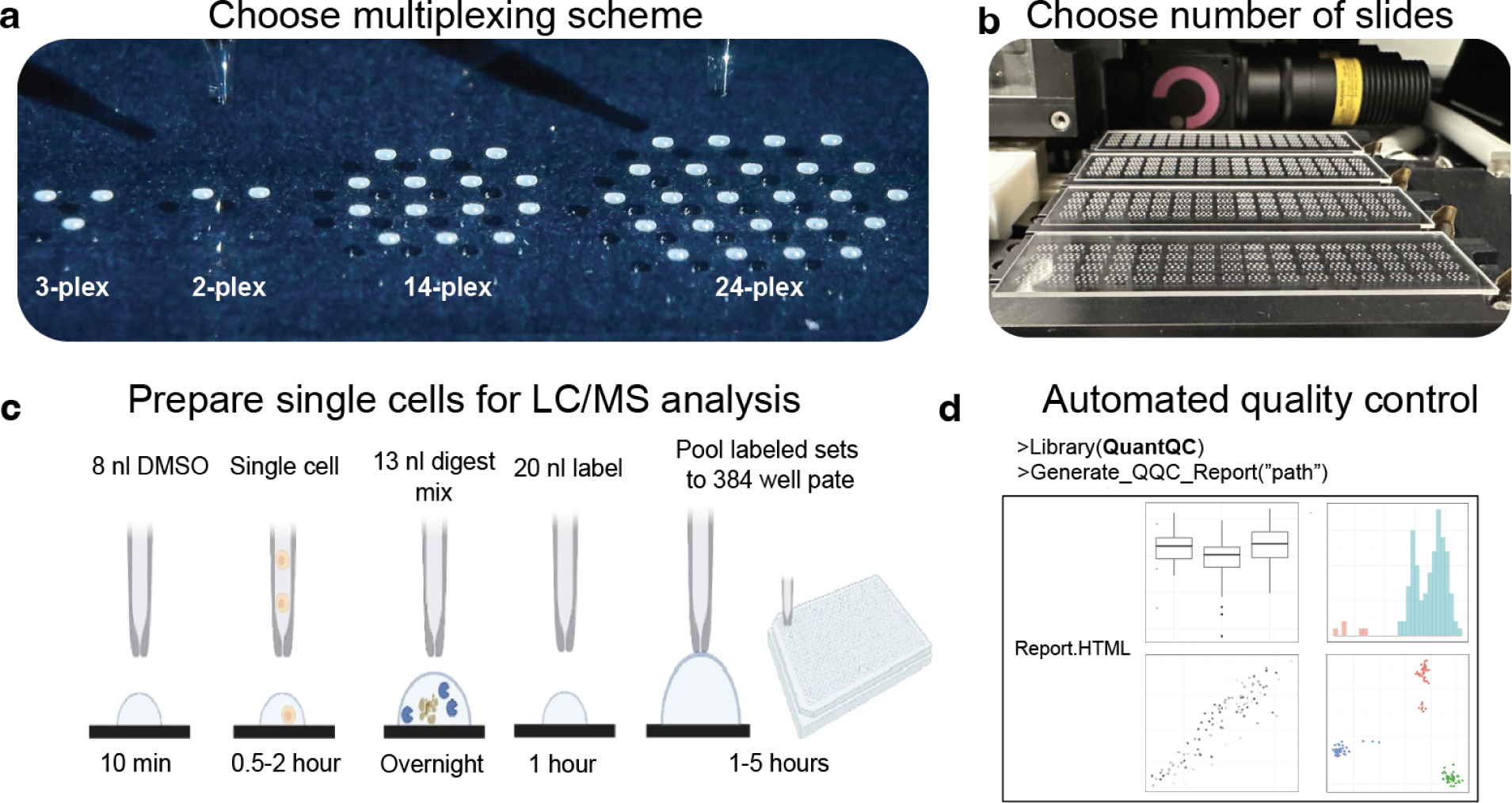
nPOP workflow. **a**, nPOP is a proteomic sample preparation method that prepares single cells in droplets on the surface of glass slides. This allows for flexible design that can fit any desired multiplexing scheme as reflected by the number of droplets per cluster. **b**, A picture of a workflow using 4 glass slides and the 14-plex design allowing for simultaneous preparation of 3,584 single cells for prioritized proteomic analysis. **c**, A schematic of the nPOP method illustrates the steps of cell lysis, protein digestion, peptide labeling, quenching of labeling reaction, and sample pooling and the transfer of the pooled samples to an autosampler plate. These steps are performed for each single cell (corresponding to a single droplet). **d**, To analyze data generated from an nPOP sample preparation, the QuantQC R package can be used to map all metadata and generate quality reports for quick evaluation of the experiment.

**Figure 2:**
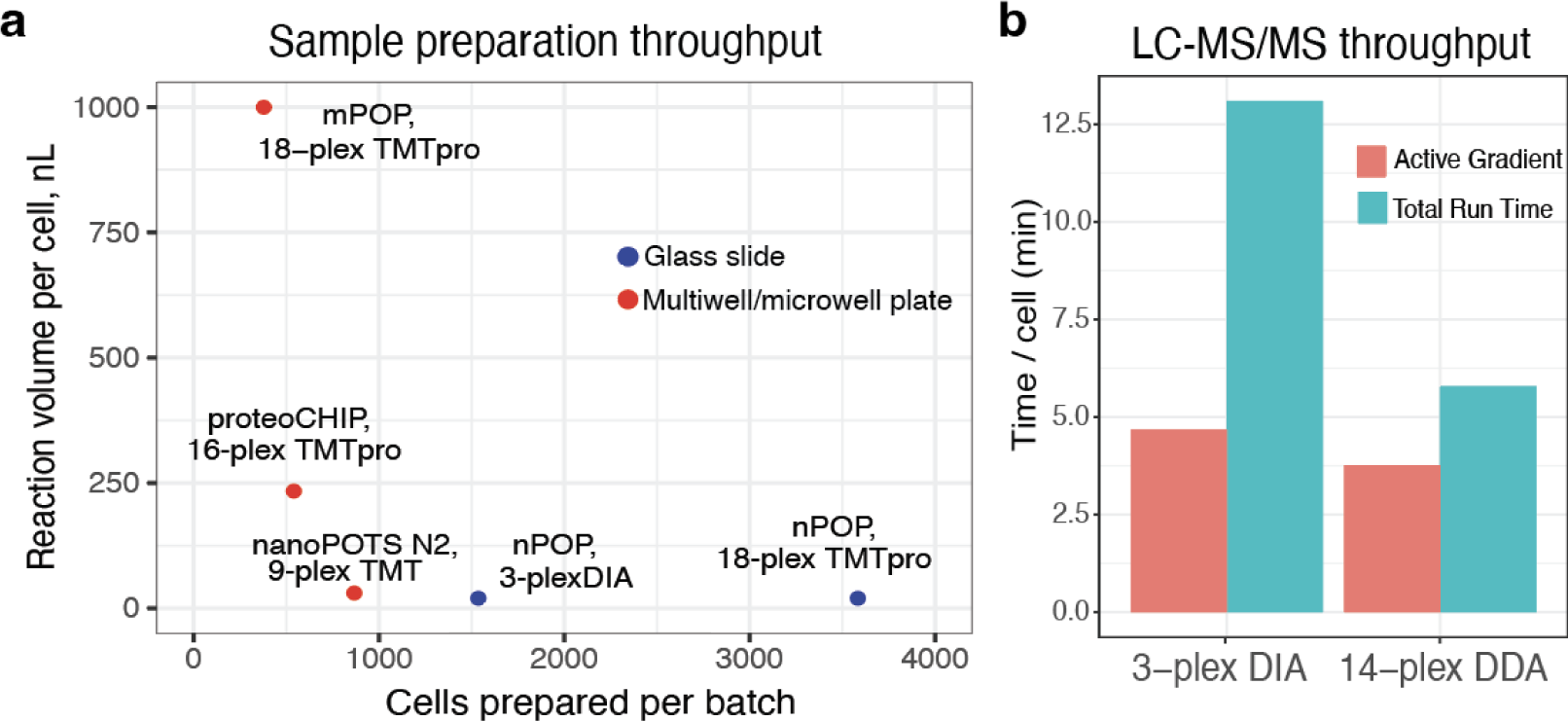
Comparison of throughput and reaction volume for multiplexed methods. **a**, Multiplexed sample preparation methods are shown the space of reaction volumes and number of cells that can be simultaneously prepared. Nanoliter reaction volumes increase efficiency of sample preparation by reducing adsorptive losses while reducing cost per cell. Slide based methods (nPOP) allow for the largest batch size, followed by microfabricated chip based methods (nanoPOTS^41^, N2^40^, proteoCHIP^12^), and multiwell plate based methods, mPOP^38,39^. **b**, Time needed for the analysis of a single cell by 3-plexDIA (about 15 min active gradient and about 37 min total run time per labeled set) and by pSCoPE (about 60 min active gradient and 85 min total run time per labeled set).

**Table 1.** Cost of sample nPOP reagents per cell. These estimates assume that the glass slide is used at full capacity as shown in Fig. 1 and account for reagents overheads detailed in this protocol.

|  |  |
| --- | --- |
| TMTpro/mTRAQ labels | \$0.08 / cell |
| Promega Trypsin Gold | \$0.01 / cell |
| TEAB | > \$0.01 / cell |
| Hydroxylamine | > \$0.01 /cell |
| H1 glass slide | \$0.04 / cell |
| TOTAL | ~ \$0.12 / cell |

nPOP allows for preparing samples utilizing multiple multiplexed workflows, including plexDIA with mTRAQ and the SCoPE experimental designs with TMT 11-plex, TMTPro 18-plex and 27-plex isobaric mass tags. The SCoPE samples can then be analyzed using prioritized data acquisition to maximize the coverage and biological relevance of the single cell data ^28^. nPOP sample preparation can easily include the use of a carrier channel or new higher plex mass tags that can substantially increase the throughput of single-cell MS proteomics ^21,26^.

The protocol described here has a few modifications of the version 1 protocol first reported by Leduc *et al.*^7^.These include obviation of the perimeter droplets (which we found not to be needed), an nPOP software module to facilitate protocol execution, an (optional) overnight protein digest, and an R package QuantQC facilitating data evaluation and initial analysis.

### Experimental design and mass tag choice

Single cells can be multiplexed using different types of mass tags, including isobaric or and non-isobaric mass tags, and their tradeoffs determine which multiplexing workflow best suits experimental needs. Currently, the TMTpro workflow provides the highest sample throughput, Fig 2b. plexDIA has lower throughput, but its compatibility with shorter LC gradients and the development of higher plex non-isobaric mass tags can substantially increase its throughput^21,26,29^. Another difference is that protein quantification using the reporter ions of isobaric mass tags is adversely affected by coisolation interferences while quantification with non-isobaric mass tags is not affected. Ultimately, using isobaric and non-isobaric approaches to cross-validate each other enables higher confidence as demonstrated by Leduc *et al.*^7^ and recommended by the community guidelines^1^.)

### Data Analysis

To facilitate robust and standardized analysis of data from nPOP experiments, we converted the analysis pipeline developed in Leduc *et al.*^7^ into the QuantQC R package. QuantQC generates HTML reports for evaluating nPOP sample preparation, stability of data acquisition, and quantification performance that can be easily shared with colleagues. QuantQC also facilitates exploratory data analysis such as visualizing agreement between peptides mapping to the same protein across clusters.

### Application of the method

nPOP has been applied to study various cell lines ^7,17,30^ as well as primary cells^28^ and tissue samples. nPOP can be applied to tissues when a suspension of whole cells can be generated similarly to droplet based single cell RNA sequencing methods. In addition to single cells, the nPOP sample preparation can be applied to small subsets of cells, clusters of interacting cells or larger whole organoids that can be isolated by the speroONE instrument that sorts larger particles. So far, nPOP has been used for bottom up proteomics, but in the future it may be adopted to increase the throughput of emerging top-down methods for analyzing intact proteoforms^31,32^.

### Comparison to other methods

Many methods can prepare single cells for label-free LC/MS analysis ^9,33–37^. nPOP can also be used for preparing cells for label-free analysis, but here we focus on its multiplexed applications and compare them to other methods that support multiplexed sample preparation since multiplexing can support higher throughput, only a few minutes per single cell as shown in Fig. 2.

Multiplexed methods for preparing many single cells can be categorized as using: (i) multiwell plates, (ii) microfabricated chips, and (iii) glass slides. The Minimal ProteOmic sample Preparation (mPOP)^38,39^ falls in the first category, a multiwell plate based sample preparation method. It has been used to prepare single cells for LC/MS analysis. mPOP is performed in a 384 well plate in volumes of about 1 microliter per cell and requires manual pooling of labeled samples prior to LC-MS/MS analysis. Other multiplexed methods for preparing single cells for protein analysis by LC/MS including the proteoCHIP^12^, nanoPOTS^40^, and its more recent version N2^40^. These methods utilize microfabricated chips to achieve small volume sample preparation. The benefits of nPOP’s glass slide based approach include the relative ease of fabricating glass slides, the spatial flexibility of being able to dispense droplets anywhere over the slides surface, and the higher throughput allowing for simultaneous preparation of thousands of single cells, Fig. 2. This not only allows nPOP to adapt to any multiplexing scheme without the need to redesign chips, it also allows nPOP to achieve the lowest volume and highest throughput sample preparation of similar methods. A comparison of the volumes utilized by these different sample preparation methods and the number of cells that can be prepared in one preparation can be found in Fig. 2.

Benchmarking efficiency of sample preparation can be challenging without access to samples prepared by alternative methods of matched cell types run on the same instrument in a similar time frame. However, nPOP has demonstrated competitive protein coverage in diluted standards ^7^. Quantitative accuracy has also been benchmarked using synthetic peptides spiked into single cell proteomes spanning a 16-fold dynamic range ^28^, which has not been demonstrated by other methods.

### Expertise needed to implement the protocol

In order to implement the nPOP sample preparation workflow, familiarity and basic training with the CellenONE system is required. Cellenion offers an nPOP Partnership Program to accelerate the successful implementation of nPOP with custom accessories, access to process experts, and the latest protocols. Executing the protocol properly may take one to two attempts, and it is advised for experimenters to not handle precious samples until they are comfortable with the workflow. Additional aspects of the workflow should be easily executable for a biochemist or cell biologist with cell culture and molecular biology experience. Familiarity with considerations of proteomic sample preparation is also advised. The full nPOP protocol can be effectively completed by a single user. Additional users may be beneficial for executing more complex experimental designs such as those that require sorting populations of cells across a plurality of treatment conditions, as is in harvesting cells from a time-course stimulation.

### Limitations

nPOP requires use of specialized equipment, the CellenONE, that limits usability of the method in labs that do not have access to this instrument. Additionally, performing the cellenONE sample preparation requires the user to be a proficient operator of the instrument, which may take up to several days of practice. These limitations are partially mitigated by the possibility of preparing samples in one laboratory and then shipping them on dry ice for analysis in another laboratory, as has been demonstrated^17^.

Limitations in the speed of cell dispensing limit the practical throughput of sample preparation to ∼3500 single cells per single prep, through modifications of CellenONE can substantially increase dispensing speeds and relax this limitation. Dispensing thousands of cells takes up to two hours, so if the user has sensitive samples, long isolation times may be suboptimal. Additionally, if the tissue sample is challenging to dissociate and only nuclei suspensions are feasible, the experimenter will only be able to measure nuclear proteins. If samples are contaminated with chemicals undermining MS analysis, nPOP will not be able to effectively remove them. Such contaminated samples may be prepared for MS analysis with other methods, such as SP3^42^ or STrap^43^.

Lastly, nPOP requires suspension cells limiting its application to frozen tissue samples that may not be amenable to generating a whole cell suspension.

## Materials

### Biological materials

- Isolated primary cells, or cell lines grown in culture. Multiple cell lines can be used for preparation of initial nPOP samples. To generate a small-scale nPOP experiment that is featured within this manuscript, we used THP1, WM989, and CPAF pancreatic cancer cell lines for plexDIA analysis.

### Reagents

- Water, Optima LC-MS/MS grade (Fisher Scientific, cat. no. W6-1)
- Acetonitrile (for buffer preparation), Optima LC-MS/MS grade (Fisher Scientific, cat. no. A955-1)
- Triethylammonium bicarbonate (TEAB), 1 M pH 8.5 (Sigma Alrich, cat. no. T7408100ML)
- Formic acid, Pierce, LC-MS/MS grade (Thermo Fisher Scientific, cat. no. 85178)
- n-Dodecyl-beta-Maltoside Detergent (Thermo Fisher Scientific, cat. no. 89902)
- TMTpro 18-plex Label Reagent Set, 1 × 5 mg (Thermo Fisher Scientific, cat. no. A44520)

OR

TMTpro 27-plex Label Reagent Set

OR

mTRAQ Reagents (Sciex, cat. no 4374771)

- Hydroxylamine (HA), 50% wt/vol (Sigma, cat. no. 467804-50ML)
- Trypsin, Trypsin Gold Mass Spectrometry Grade (Promega, cat. no. V5280)
- Phosphate-buffered saline (PBS), 10X, pH 7.4, RNase-free (Thermo Fisher Scientific, cat. no. AM9625

### Equipment

- CellenONE cell dispenser and liquid handling robot (Cellenion)
- SciCHIP H1 coated glass slides (Scienion, cat no. CSC-5325-25)
- CellenVIALs (Scienion, cat no. CEV-5801-500)
- PCR plate, 384-well, standard (Thermo Fisher Scientific, cat. no. AB1384)
- Adhesive PCR plate foils (Thermo Fisher Scientific, cat. no: AB0626)
- PCR tubes: TempAssure 0.2 mL PCR 8-tube strips (USA Scientific, cat. no. 1402-3900)
- Plate spinner, e.g., PlateFuge microcentrifuge (Benchmark Scientific, Model C2000). This plate spinner does not offer speed control as it is used to collect liquid at the bottom of a well, rather than for pelleting material
- SpeedVac that can dry down 384 well plates on low heat or lyophilize
- **(Optional)** Mantis microfluidic liquid handler (Formulatrix)
- **(Optional)** MANTIS Chip – Silicone, HV (1μL & 5μL) (Formulatrix, cat. no. 233580)
- **(Optional)** Mantis PCR plate adapter with wide conical pins for automated plate handling (Formulatrix, cat. no. 232400)

### Software

- nPOP module for the CellenONE Software (Cellenion) provided with the nPOP Partnership Program. Earlier versions of the protocol ^7,44^ may be performed without this module, but this module is very helpful and strongly recommended.
- A DDA search engine, such as MaxQuant software (v2.4.2 or newer), available at https://www.maxquant.org with free registration or FragPipe^45^, or other DDA search engines.
- A DIA search engine, such as DIA-NN software (v1.8.1 or newer)^46^, available at https://github.com/vdemichev/DiaNN/releases/tag/1.8.1, Spectronaut^47^, or MaxDIA^48^, available at https://www.maxquant.org.
- **(Optional)** Software for rescoring peptides identified by the search engines by including retention time information (e.g., DART-ID^49^) or peptide fragmentation patterns (e.g., MSBooster^50^, Oktoberfest^51^), and other features (e.g., Mokapot^52^).
- **(Optional)** QuantQC R package for quality control, available at https://scp.slavovlab.net/QuantQC and DO-MS for optimizing LC-MS/MS data acquisition parameters available at: https://do-ms.slavovlab.net/ and https://github.com/SlavovLab/DO-MS
- **(Optional)** Pipelines for data processing including the scp R–Bioconductor package^53,54^ and the SPP Pipelines ^55^ available at https://github.com/SlavovLab/SPP.

## Procedure

### Software connection and system initialization

The current version of the nPOP protocol described in this article requires software provided by the manufacturer of the CellenONE X1 system.

1. Initialize the instrument with the “nPOP” user folder.
2. Ensure that the instrument chiller is powered on.
3. Prime the instrument with nozzles in position 1 and 3 by following the on-screen prompts.

a. For optimal pickup recovery, a difference of <50 μm in the z-offset between nozzles should be observed
4. Fill DI water to the fill line in the cellenONE humidifier

Once initialized and primed, each step of the protocol is preprogrammed with automatic environmental controls and suggested sample conditioning via on-screen prompts.

### Preparing cells for sorting and bulk injection

Cells are needed for two purposes, a) sorting single cells for the nPOP experiment and b) to make bulk samples for either carrier or DIA library generation. Cells can be obtained fresh from culture, or from a dissociated cell suspension frozen at −80 or liquid nitrogen in a solution of 10% DMSO 90% 1x PBS.

For cell sorting, cells should be washed of media / cryopreservative and resuspended in 1x PBS at a concentration of 300 cells per microliter. For bulk samples, cells should be suspended at 2000 cells per μL in LC/MS grade water and frozen at −80 for future use.

### nPOP Procedure

nPOP consists of 6 required steps depending on the multiplexing scheme designed for the single-cell experiment. An optional first step can be taken to spike synthetic peptides into single cell proteomes. Should the user not desire to utilize this feature of nPOP then they may proceed with all subsequent steps as described. Importantly, on-screen prompts following the initiation of each step allow the user to validate that optimal system functionality is achieved before proceeding.

The user can decide how many glass slides of cells they would like to prepare corresponding to the number of cells they would like to ultimately analyze. This choice can be carried out by minimizing the number of fields in the y-axis. Droplet patterns are repeated in units called fields. Each field has dedicated quality control spots and each slide has 4 fields for a total of 16 fields. The user can also dispense to a quarter, half, or three quarters of a slide if desired, but this is not recommended.

Choices for different slide numbers primarily change the time required for cell sorting and for sample pooling. Considerations for the number of slides to prepare are listed table 2 below:

**Table 2.** Slide number considerations. The number of slides and multiplexing scheme chosen will dictate the number of single cells prepared in one experiment.

| Multiplexing scheme | Number of slides | Number of cells prepared |
| --- | --- | --- |
| TMTpro 14-plex | 1 | 896 |
|  | 2 | 1792 |
|  | 3 | 2688 |
|  | 4 | 3584 |
| mTRAQ 2-plex | 1 | 320 |
|  | 2 | 640 |
|  | 3 | 960 |
|  | 4 | 1280 |
| mTRAQ 3-plex | 1 | 384 |
|  | 2 | 768 |
|  | 3 | 1152 |
|  | 4 | 1536 |

### Part 0: Select multiplexing workflow

The following protocol describes preparing samples for analysis by 3-plex plexDIA, 2-plex plexDIA allowing for the use of a carrier, and 14-plex TMT for DDA or prioritized data acquisition. New multiplexing protocols will be provided once validated and can be requested through the nPOP Partnership Program.

### Part 1: Synthetic peptide spike-in (optional)

1. Prepare a solution of synthetic peptides in LC-MS grade water such that the amount per 300 picoliters represents the lowest desired level of peptide copies across the dynamic range.
2. In the main tab, set the run to: “0_Dispense_SpikeIns”.
3. CRITICAL: When dispensing synthetic peptide spike-ins, the cooling and humidity control should be turned OFF. This will allow the synthetic peptides to dry onto the glass slide, which will be resuspended when the volume of lysis reagent is dispensed.
4. Under Target Setup, Load the SpikeIns.fld field file from the relevant multiplexing scheme folder and set the No. of Fields “Y” to match the number of slides intended for processing.
5. Load 150 μL of the synthetic peptide stock solution into a fresh CellenVIAL and place into position 2 of the CellenWASH station with the cap pointed towards the door of the system.
6. Select the “Run” tab and click “Start Run”. Follow on-screen prompts. Once dispensing is complete, the nozzles will be flushed and images will be taken across the glass slides for visual inspection of the quality and distribution of droplets.

### Part 2: Dispense DMSO

6. In the main tab, set the run to “1_Dispense_DMSO”.
7. Under Target Setup, Load the DMSO.fld field file from the relevant multiplexing scheme folder and set the No. of Fields “Y” to match the number of slides intended for processing.
8. Load 150 μL of LC-MS grade DMSO into a fresh CellenVIAL and place into position 2 of the CellenWASH station with the cap pointed towards the door of the system.
9. Under the Run tab, select Start Run. Follow on-screen prompts.
a. The first step will aspirate 10 μL of DMSO and ask the user to manually confirm the stability of the droplet. Test droplet with autodrop 3-5 times. If droplet is consistent throughout 3 straight droplets, hit Continue Run on the pop up menu.

TROUBLESHOOTING

10. Once dispensing is complete, the nozzles will be flushed and images will be taken across the glass slides for visual inspection of the quality and distribution of droplets. CRITICAL: Check images of DMSO droplets to ensure uniform dispensing over all slides. Check for any missing droplets or signs of off target dispense.

TROUBLESHOOTING.

### Part 3: Dispense Cells

11. Prepare cell populations within 1 hr of sorting by first washing twice in a suspension of 1x PBS and store on ice until loaded into the instrument for sorting. Aim for a final concentration of 200-300 cells per microliter. Counting cells by hemocytometer is recommended.

CRITICAL: If cell suspension is prone to aggregation, run through a 40 micron strainer prior to aspiration to avoid nozzle clogging. If cells larger than 40 micron are of interest and likely present in a sample, a 70 micron strainer can be used instead to remove very large aggregates.

12. In the main tab, set the run to “2_Dispense_Cells”.
13. Under Target Setup, navigate to the CellsFieldFiles folder in the relevant multiplexing scheme folder and set the No. of Fields “Y” to match the number of slides intended for processing.
a. Field files are provided for if the experiment requires 1 or 2 conditions. If using one condition, navigate to the 1_condition folder and load the cells.fld field file. If multiple conditions, select CellType_A.fld or CellType_B.fld field files.
14. Load at least 50 μL of cell suspension into a fresh CellenVIAL and place into position 1 of the CellenWASH station with the cap pointed towards the door of the system.
a. For samples with a limited number of cells (<10,000) 30 uL can be loaded into a 384 well PCR plate to decrease dead volume and recover the maximum number of cells for processing.
15. Under the Nozzle Setup tab in the Do Task menu, perform the Take10ul_CellenVIAL task to aspirate cell suspension.
a. For sample aspiration from the 384 well PCR plate, use the take probe hot button and leave at least 5 uL dead volume in the well to avoid aspirating air.
16. Under the Nozzle Setup tab, open the CellenONE cell dispensing window and run the mapping task to set the ejection zone and identify the cells of interest from the size and elongation distribution.
a. CRITICAL: You may want to adjust the cell size and elongation distribution manually based on the goals of your experiment.
17. When parameters are sufficiently tuned, under Run, select Start Run. Follow on-screen prompts. Once dispensing is complete, the nozzles will be flushed and images will be taken across the glass slides for visual inspection of the quality and distribution of droplets.
a. CRITICAL: If dispensing thousands of cells, it is useful to periodically pause the run by pressing the nozzle setup button to assure the droplet is still stable.

TROUBLESHOOTING.

18. Repeat steps 14-22 for any additional cell suspensions prepared for sorting.
19. Once finished sorting the cells, close the CellenONE cell sorting operation window.
20. Store remaining cell suspensions on ice until the end of Part 3 of the protocol. The remaining cell suspensions can be used to generate bulk samples for empirical library building, additional analysis, and validation. If not needed, any remaining cell suspensions can be disposed of in appropriate waste containers.

### Part 4: Dispense Digest Master Mix

21. In the main tab, set the run to “3_Dispense_Digest”.
22. Under Target Setup, Load the Digest.fld field file from the relevant multiplexing scheme folder and set the No. of Fields “Y” to match the number of slides intended for processing.
23. Prepare a fresh Digest Master Mix stock solution of 100 ng/μL Trypsin Gold, 5 mM HEPES, and 0.05% DDM in LC-MS grade water.
24. Load 150 μL of Digest Master Mix into a fresh CellenVIAL and place into position 2 of the CellenWASH station with the cap pointed towards the door of the system.
25. Under the Run tab, select Start Run. Follow on-screen prompts. Once dispensing is complete, the nozzles will be flushed and images will be taken across the glass slides for visual inspection of the quality and distribution of droplets.
a. Inspect droplet images to ensure Digest Master Mix was added to each droplet successfully.

TROUBLESHOOTING

26. Allow the proteins to digest within each droplet for at least 3 hours, ideally 8 hours or overnight.

TROUBLESHOOTING

PAUSE POINT:

End of day one. Beginning day two, proceed with mTRAQ labeling (Part 4) or TMT labeling (Part 5) according to the chosen experimental design for either plexDIA or pSCoPE, respectively.

### Part 5: Dispense Labels

27. Prepare stock solutions of mTRAQ or TMTpro for labeling single cells for multiplexed analysis.

For mTRAQ, transfer 10 μL of stock concentration for each mTRAQ label to a PCR tube repeating each label for each slide being prepared. For example, if the user is preparing 4 slides of sample for 3plex analysis, they will have 12 PCR tubes each with 10 μL of label, 4 of mTRAQ d0, 4 of mTRAQ d4, and 4 of mTRAQ d8.

OR

For TMTpro, transfer 10 μL of each label into a PCR tube.

28. Dry down each tube in a SpeedVac vacuum concentrator on the low heat setting. Drying should take up to 10-12 minutes.
29. Resuspend the tags in the PCR tubes with LC-MS grade DMSO. Pippet mix well to ensure proper redissolving of the labels.

mTRAQ tags will be resuspended in 20 μL per tube for a 2X dilution from the stock concentration.

OR

TMT tags will be resuspended in 30 μL per tube for a 3X dilution from the stock concentration.

30. Load 20 μL of mTRAQ tag, or 30 μL of TMT tag into the wells of a 384-well plate to be placed inside the CellenONE X1 system. Load labels beginning in position G1, as follows:
31. In the main tab, set the run to “4_Dispense_Labels”.
32. Under Target Setup, Load the Labels.fld field file from the relevant multiplexing scheme folder and set the No. of Fields “Y” to match the number of slides intended for processing.
33. Under the Run tab, select Start Run. Follow on-screen prompts. Once dispensing is complete, the nozzles will be flushed and images will be taken across the glass slides for visual inspection of the quality and distribution of droplets.

TROUBLESHOOTING

34. If using mTRAQ labels, immediately after dispensing labels, proceed to Dispense TEAB (Part 5) of the procedure. If using TMT labels, let labels incubate for 1 hour after the last label is dispensed.

### Part 5: Dispense TEAB (mTRAQ only)

35. Prepare TEAB solution for dispensing immediately after mTRAQ dispensing in Part 4. The TEAB solution should be prepared fresh to 100 mM in LC-MS grade water and verified with pH of pH 8.0-8.4.
a. NOTE: this step may be completed during the dry-down of the mTRAQ stock solution listed above in step 35.
36. Load 150 μL of 100 mM TEAB solution into a fresh CellenVIAL and place into position 2 of the CellenWASH station with the cap pointed towards the door of the system.
37. In the main tab, set the run to “5_Dispense_TEAB”.
38. Under Target Setup, Load the TEAB.fld field file from the relevant multiplexing scheme folder and set the No. of Fields “Y” to match the number of slides intended for processing.
39. Under the Run tab, select Start Run. Follow on-screen prompts. Once dispensing is complete, the nozzles will be flushed and images will be taken across the glass slides for visual inspection of the quality and distribution of droplets.

TROUBLESHOOTING

40. Once TEAB dispensing has been completed, allow labels in the buffer to incubate for 1 hour, then proceed to Sample Pickup (Part 8) of the procedure.

### Part 6: Dispense HA (TMT only)

41. Prepare hydroxylamine (HA) solution for dispensing after TMT labeling incubation in Part 5. The HA solution should be prepared fresh to 5% in LC-MS grade water.
42. Load 150 μL of 1% by weight solution of Hydroxylamine into a fresh CellenVIAL and place into position 2 of the CellenWASH station with the cap pointed towards the door of the system.
43. In the main tab, set the run to “6_Dispense_HA”.
44. Under Target Setup, Load the HA.fld field file from the relevant multiplexing scheme folder and set the No. of Fields “Y” to match the number of slides intended for processing.
45. Under the Run tab, select Start Run. Follow on-screen prompts. Once dispensing is complete, the nozzles will be flushed and images will be taken across the glass slides for visual inspection of the quality and distribution of droplets.
a. For droplets that appear to have had issues with HA dispensing, the user can repeat labeling steps 55-57 to ensure enough buffer has been introduced to each droplet. Ensure enough HA solution remains for use inside the CellenVIAL. Be cautious to not over dispense HA volumes, as introducing the lowest concentration possible into samples typically results in better peptide sequence identification during LC-MS/MS.
46. Once HA dispensing has been completed, allow labels to quench for 30 minutes, then proceed to Sample Pickup (Part 8) of the procedure.

### Part 7: Sample Pickup

47. Prepare a fresh stock of sample pickup solution. Load 10 mL of 50:50 acetonitrile:water and 0.1% formic acid (LC-MS grades) into the CellenONE WashTray XL.
a. IMPORTANT: when placing the WashStation into the service station slot be careful not to touch the glass slides and mounted PDC nozzles.
48. Prepare one (if TMT or 1-2 slides prepared with plexDIA) or two (if 3-4 slides prepared with plexIDA) fresh 384-well plates that contain 2 μL of 0.01% DDM in water (LC-MS grade) in each well. Place the first plate inside the CellenONE. Store the second plate in a 4°C until the second pickup step is reached.
a. NOTE: label the plates according to order of sample pickup. For plexDIA the first two glass slides will be placed into the first plate via automatic pickup by the CellenONE system. The last two glass slides will be placed into the second plate, after the first round of sample pickup is complete.
49. In the main tab, set the run to either “7_Pickup_plexDIA” or “7_Pickup_TMT” depending on the relevant workflow.
50. Under Target Setup, Load the Pickup.fld field file from the relevant multiplexing scheme folder and set the No. of Fields “Y” to match the number of slides intended for processing.
51. Under the Run tab, select Start Run. Follow on-screen prompts. Once sample pickup is complete, the nozzles will be flushed and images will be taken across the glass slides for visual inspection of the glass slides to ensure all samples have been picked up. When preparing thousands of single cells, this process may take approximately up to 2 hours for TMT workflow and 6 hours for the 2-plex workflow.
52. Review the pickup images to confirm limited residual reaction volumes are present.

TROUBLESHOOTING

53. Once samples from slides one and two have been picked up and loaded into the first 384-well plate, seal the plate with an adhesive foil plate cover, spin down to collect the droplets in the bottom of the wells
a. Once collected, remove the foil to begin drying down the samples on low heat in a SpeedVac vacuum concentrator. This process may take up to an hour. Store the plate at −80°C until ready for resuspension and LC-MS/MS analysis.
b. NOTE: while drying down, allow the third and fourth slides to begin the sample pickup step.
54. Load the second plate into the CellenONE system to begin the final round of pickup. Repeat steps 70-74 for the second plate, following through with drying down the plate and storing according to the conditions in step 74.

### System Cleaning and Shutdown

#### CellenONE shutdown

1. Once the final 384-well plate containing prepared single-cells is drying down, the user may begin the system cleaning and shutdown procedure.
2. Turn off the humidity and temperature control. Carefully remove the humidity control tubing and rinse with 70% IPA or EtOH and allow to dry prior prior to reinstallation before subsequent experiments
3. Replace the acetonitrile:water solution in the CellenOne WashStation with 100% Isopropyl alcohol (LC-MS grade) and place it back inside the system.
4. Under the DoTask menu in the Nozzle Setup tab, run the nPOP_EndOfRun_Wash.

a. The nozzles will move inside the pool of isopropyl alcohol and soak for four hours. The user may return later to complete shutdown, or allow the system to complete this step overnight.
5. Once all steps are complete, close the CellenONE software. Turn off the computer by selecting “shutdown”. Once the computer is fully shut down, then turn off the power to the CellenONE X1 system.

#### Manual nozzle cleaning

For optimal cell sorting and reagent dispensing, thoroughly clean PDCs are required every 2-3 sample preparations.

1. Carefully remove PDC from CellenONE X1 system according to manufacturer’s instructions.
2. Connect a 20 mL Luer lock plastic syringe to the end of the PDC tubing, carefully leaving an air gap of approximately 5mL.
3. Fill a 100 mL glass beaker with distilled water and place it into a water bath sonicator.
4. While sonicating, place the glass tip of the PDC connected to the syringe into the distilled water inside the beaker, then carefully withdraw water to begin flowing through the PDC, the plastic tubing, and into the syringe. Allow a large droplet of water to form inside the plastic syringe. The aim is to withdraw any large particles of debris potentially stuck inside the PDC into the syringe to be disposed of without exiting the small diameter of the glass tip.

a. CRITICAL: allow only the glass tip of the PDC to be exposed to water, otherwise potential damage and loss of performance may arise due to short circuit of the piezo ceramic
5. Disconnect the plastic syringe, and dispose of the water and potential debris into waste.
6. Reconnect the PDC and syringe with a 5 mL air gap via Luer lock.
7. While sonicating, place the glass tip of the PDC back into the distilled water in the beaker, and begin withdrawing water into the PDC for 5-8 seconds. Withdraw enough water to fill the PDC itself, but not filling the tubing and syringe of the connection. Filling beyond the volume of the PDC is unnecessary.

a. CRITICAL: be careful not to touch the glass tip of the PDC to the walls of the beaker, as the tip may break, coating may be removed, or become otherwise compromised.
8. Carefully push down on the syringe while keeping the glass tip of the PDC inside the beaker of distilled water. This will partially clean the PDC tip physically by using slight force and sonication. Once all the withdrawn liquid has been ejected, a steady stream of bubbles should be observed inside the beaker of distilled water.
9. Repeat steps 7 and 8, several times. After each repeat of step 8, check the spray of the PDC tip by removing the tip from the distilled water while applying pressure. A constant, steady, and straight stream should be observed. After several cycles, ensure all the water has been dispensed by observing another steady stream of bubbles when dispensing while the tip of the PDC is inside the distilled water.
10. Prepare a 50 mL plastic conical tube by filling with 30-50 mL of LC-MS grade ethanol.
11. While maintaining the approximately 5 mL air gap in the syringe, carefully withdraw ethanol into the empty PDC for another 5-8 seconds. Ensure the ethanol is withdrawn only inside the PDC and does not reach the tubing connection with the syringe.

a. CRITICAL: be careful not to touch the glass tip of the PDC to the walls of the beaker, as the tip may break, fracture, or become otherwise compromised. Allow only the glass tip of the PDC to be exposed to ethanol, otherwise potential damage and loss of performance may arise with exposure to the metal components of the PDC.
12. Carefully push down on the syringe while keeping the glass tip of the PDC inside the tube of ethanol. Once all the withdrawn liquid has been ejected, a steady stream of bubbles should be observed inside the ethanol. This step further removes potential debris and contaminants from the PDC.
13. Repeat steps 11 and 12 several times. After each repeat of step 12, check the spray of the PDC tip by removing the top from the ethanol while applying pressure. A constant, steady stream should be observed. After several cycles, ensure all the ethanol has been dispensed by observing another steady stream of bubbles when dispensing while the tip of the PDC is inside the ethanol.
14. Detach the syringe from the tubing connected to the PDC.
15. Fold a small Kimwipe such that it provides a long, flat edge. Dip this folded Kimwipe into the ethanol such that the material is fully soaked.
16. Carefully and delicately use the flat edge of the ethanol-soaked Kimwipe to gently wipe only the glass tip of the PDC. This final step ensures sufficient physical clearing of potential debris on the glass tip of the PDC that previous cleaning steps did not achieve.

a. CAUTION: use extremely gentle force when wiping the glass tip of the PDC to prevent potential breakage.
17. The PDC is now sufficiently cleaned and prepared for optimal cell sorting and reagent dispensing for single-cell proteomic sample preparation and can be stored dry until next use.

### LC/MS Setup

Optimization of LC/MS setup for single-cell protein analysis has been discussed extensively elsewhere ^39,56,57^. Describing our setup briefly, TMT multiplexed data was run on an Exploris 480 mass spectrometer with the Neo Vanquish LC system and 25 cm X 75 μm ID IonOpticks Aurora separation column. mTRAQ multiplexed samples were run on timsTOF Ultra mass spectrometer with a nanoElute2 HPLC pump. The analytical column used was a 25 cm X 75 μm ID IonOpticks Aurora column with a captive spray fitting.

The plexDIA data acquisition was performed using a method with 100ms fill times and 8 PASEF frames per duty cycle, with an additional MS1 frame after every 2 MS2 frames to improve the MS1 duty cycle^17,26,57^. The LC gradient on the NanoElute2 ramped from 4 to 38% buffer B and peptides eluted for 20 minutes, with a 45 minute total run time.

### Searching MS Data

Raw LC-MS/MS data from nPOP samples using TMTpro reagents and analyzed via pSCoPE or SCoPE2 should be searched following instructions outlined in Huffman *et al*.^28^ and Petelski *et al.*^39^. Searching raw plexDIA using mTRAQ 3-plex or 2-plex should be analyzed following instructions outlined in Derks *et al.*^17^.

### Data analysis with QuantQC

Here, we describe analysis of nPOP data with the R package QuantQC, which is not required but can greatly facilitate the analysis of nPOP samples by mapping all relevant metadata from nPOP experiments, including image data collected by the CellenONE, and generating quick HTML reports for evaluating data quality. A full tutorial of the workflow can be found in supplementary file 1. Below is an overview of the analysis supported by the QuantQC package.

#### Mapping metadata

- QuantQC enables mapping CellenONE data files to the searched data. This assigns sort identities to each cell if multiple conditions were prepared or if negative controls are used to evaluate background signal, along with the diameters of the isolated cells

Required files:

Search results
Cell sorting isolation files
Csv file with 3 columns “Run”, “Well” and “Plate” linking the MS run name to the inject well of the 384 plate

#### Monitoring LC/MS performance

- Changes in LC/MS performance over the course of an experiment can lead to unwanted batch effects. QuantQC facilitates easy visualization of trends in:
- Number of precursor identifications
- MS1 precursor intensities
- MS2 fragment intensities
- Average retention time drift of precursors
- Standard deviation in retention time of precursors

#### Sample preparation quality control

- Intensity of single cells compared to the intensity of a reference or a carrier allows for calculating efficiency of peptide recovery^7^
- Digestion efficiency to monitor for incomplete digestion
- Correlation between cell volume and total protein concentration for evaluating the consistency of sample preparation
- Spiked-in peptides for benchmarking quantification accuracy as demonstrated by Huffman *et al.*^28^

#### Data processing / statistics

- Different options for collapsing peptides to protein
- Median relative peptide abundance
- DirectLFQ
- RefQuant
- Distribution of peptide and protein numbers
- Data completeness for each protein across cells and each cell across all proteins identified across runs

#### Batch effect identification

- QuantQC facilitates plotting PCA dimensionality reduction color coded by a variety of different factors that could result in batch effects such as:
- Label
- Sample type
- Summed MS signal per cell
- Run order

#### Quick report generation

- Quick generation reports allow for generating sharable PDFs with one line of code

#### Biological analysis

Clustering
Comparing consistency of peptide abundance across clusters

## Troubleshooting

Multiple quality control steps are included in each run that can allow real time identification of preparation issues and also allow for recovery from many of the common issues. This quality control is enabled by the use of 2 cameras, the Drop Camera and Head Camera.

Evaluating stability of droplets via the drop camera is an important part of using the cellenONE system. At each reagent dispensing step, the user is prompted with decision points to confirm the stability of the given reagent. In Fig. 3a,b, suitable droplets are shown for aqueous and DMSO solutions. Any droplet that deviates from these images, such as the droplet shown in Fig. 3c, may be prone to a dispense failure once reagents are dispensed to the slide. If droplet shape appears irregular, the user can first try and modulate the voltage and pulse dispense settings. If the irregularity persists, the user can opt to stop the run and should flush out the reagent and repeat the step.

**Figure 3:**
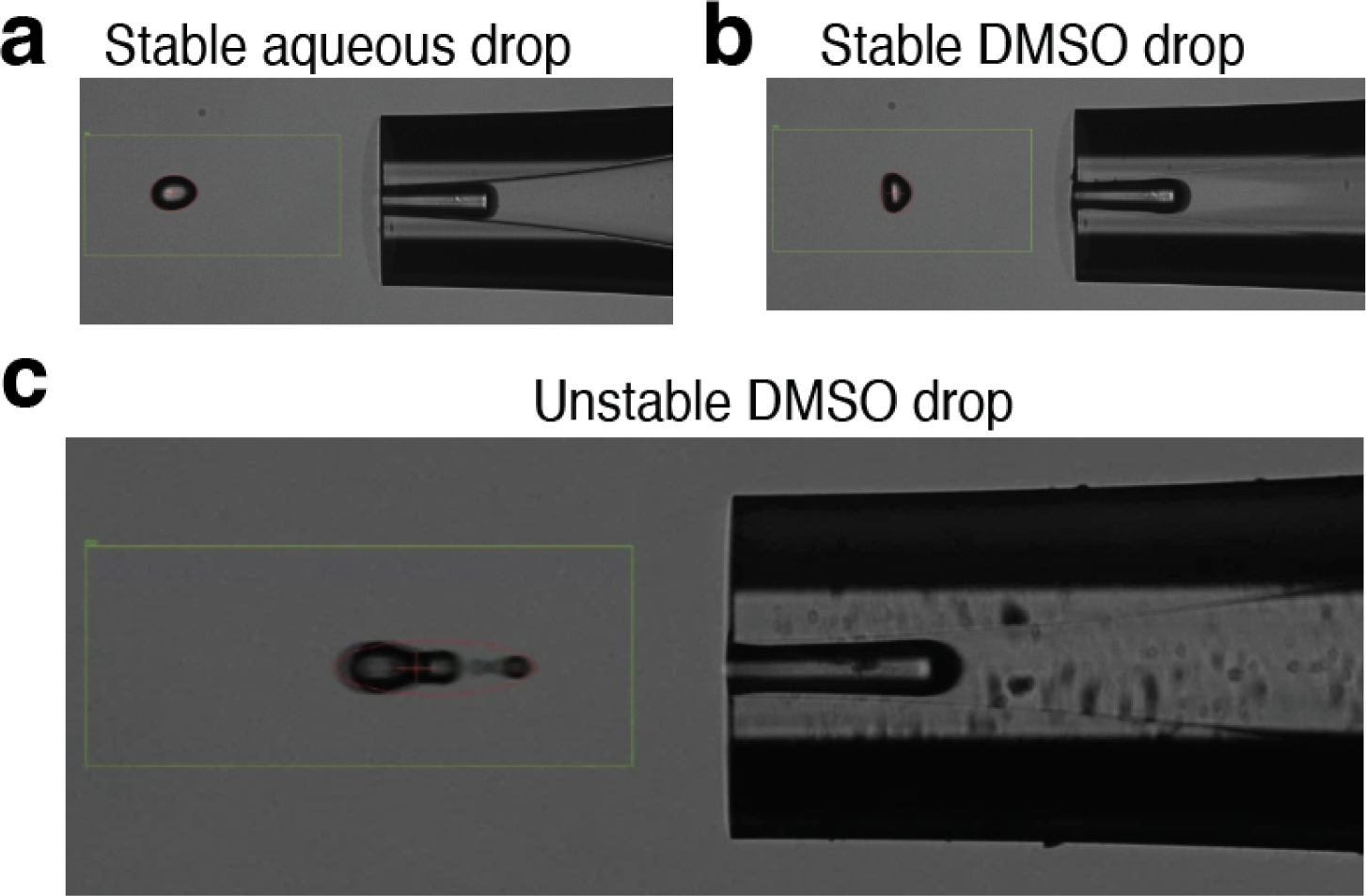
Droplet camera assessment of droplet stability. **a**, Acceptable stable droplet of an aqueous solution including cell suspensions, digest mix, TEAB buffer, and Hydroxylamine. **b**, Acceptablel stable droplet of DMSO solution including label mixtures. **c**, Possible poor droplet of DMSO showing satellite droplet requiring adjustment of voltage, pulse width, or cleaning to obtain acceptable droplet.

Head camera images allow the user to review pictures of each field on the slide after the reagents have been dispensed. In each field, all reagents are dispensed to a distinct location for the purpose of quality control, Fig 4a. If this droplet with appropriate drying characteristics is present at the bottom of the field, it suggests that reagent was prepared properly and that dispensing was successful through that field. Note, some sample preparation issues including improper pH and Trypsin activity are not assured by presence of appropriate quality control droplet features. Furthermore, examining pictures of each field can help identify where reagents should be re-dispensed Fig. 4b, or if dispensing was successful, Fig. 4c.

**Figure 4:**
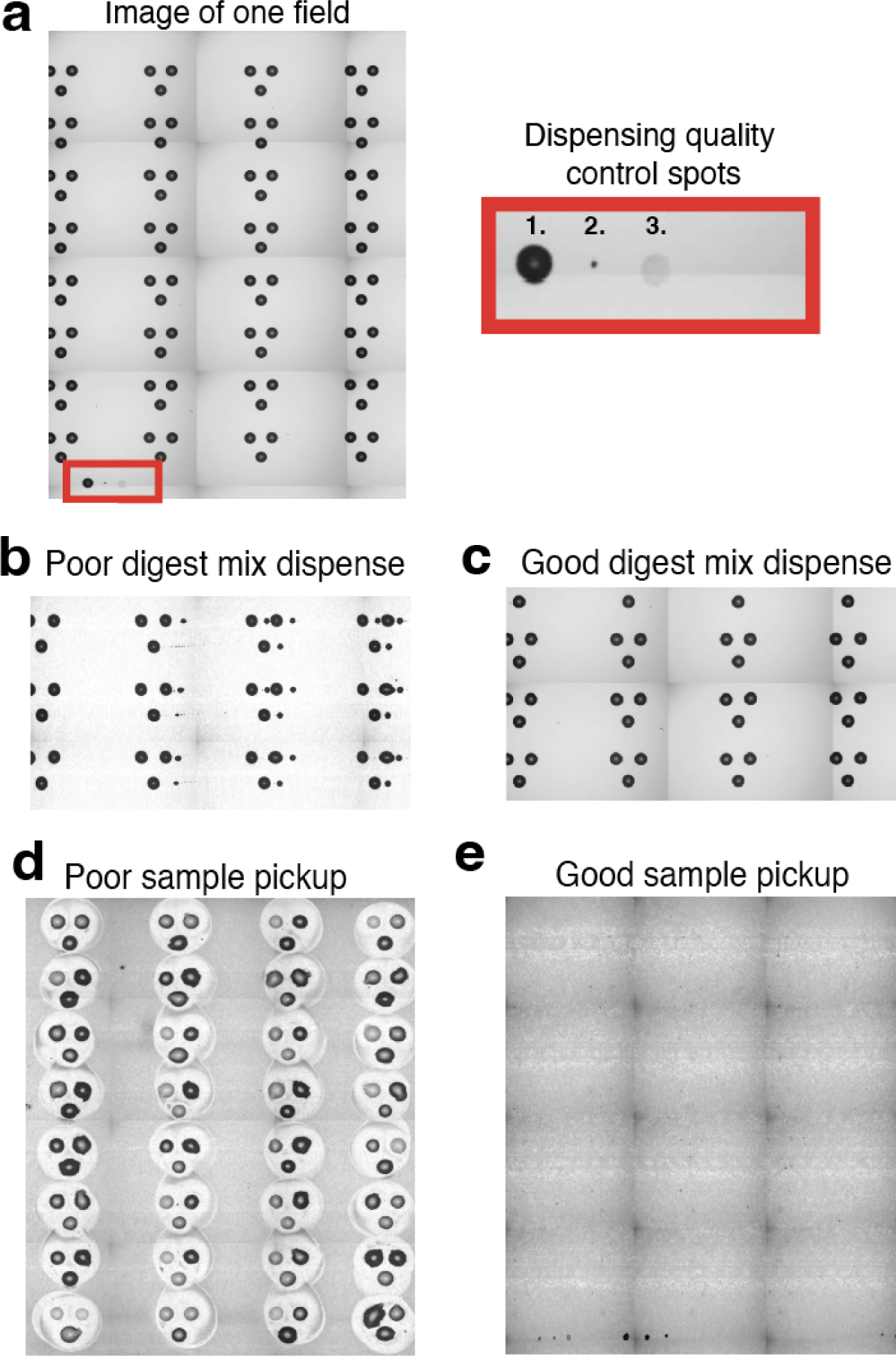
Evaluating head camera images of slides. **a**, A head camera image taken of a single field. Each slide contains 4 replicate fields. Each field contains a quality spot **1**. DMSO, **2.** Cell, **3.** Digest mix, that indicates if reagent dispensing issue occurred mid run. **b,** The user may also be able to identify failed dispense if the reagent misses the droplets. **c,** A successful dispense is often indicated by a consistent increase in the size of the reaction droplet from pre-dispense images to post-dispense images and should not show smaller droplets to the side of primary spots. **d,** An after image of failed pickup shows significant residue left behind on slide. **e,** An after image of a successful pickup shows little to no residue left on slide.

The final stage of the experiment, the sample pooling and pickup, may require some optimization when getting started with nPOP. The first important feature to optimize is the nozzle height from the slide. Poor recovery of droplets, Fig. 4d, suggests nozzles are too close or far away from the slide. This can be nozzle specific where nozzle length difference is too great or general to both nozzles in the case of improper target position point. Reach out to Cellenion for support in adjusting this parameter. Once it is at an optimal pickup height, the slide should appear as the picture in Fig. 4e after sample pickup.

Occasionally, troubleshooting steps require the user to manually edit the field files to remove unneeded droplets. To delete parts of field files that are no longer needed, after loading the field file, hold down control and select all field files that are not needed. Then go into the Field Setup tab, and erase all droplets.

Additional troubleshooting advice for individual steps can be found in Table 4.

**Table 3.** Label loadings into probe plate. The number of slides and multiplexing scheme chosen will dictate the number of single cells prepared in one experiment. If less than 4 slides are to be processed in a plexDIA experiment, only load wells correlating to the number of slides processed.

| Well position | 2-plex (mTRAQ 20 µL) | 3-plex (mTRAQ 20 µL) | 14-plex (TMT 30 µL) |
| --- | --- | --- | --- |
| G1 | d0 (Slide 1) | d0 (Slide 1) | TMTpro-128C |
| G2 | d4 (Slide 1) | d4 (Slide 1) | TMTpro-129N |
| G3 | d0 (Slide 2) | d8 (Slide 1) | TMTpro-129C |
| G4 | d4 (Slide 2) | d0 (Slide 2) | TMTpro-130N |
| G5 | d0 (Slide 3) | d4 (Slide 2) | TMTpro-130C |
| G6 | d4 (Slide 3) | d8 (Slide 2) | TMTpro-131N |
| G7 | d0 (Slide 4) | d0 (Slide 3) | TMTpro-131C |
| G8 | d4 (Slide 4) | d4 (Slide 3) | TMTpro-132N |
| G9 | - | d8 (Slide 3) | TMTpro-132C |
| G10 | - | d0 (Slide 4) | TMTpro-133N |
| G11 | - | d4 (Slide 4) | TMTpro-133C |
| G12 | - | d8 (Slide 4) | TMTpro-134N |
| G13 | - | - | TMTpro-134C |
| G14 | - | - | TMTpro-135N |

**Table 4.**
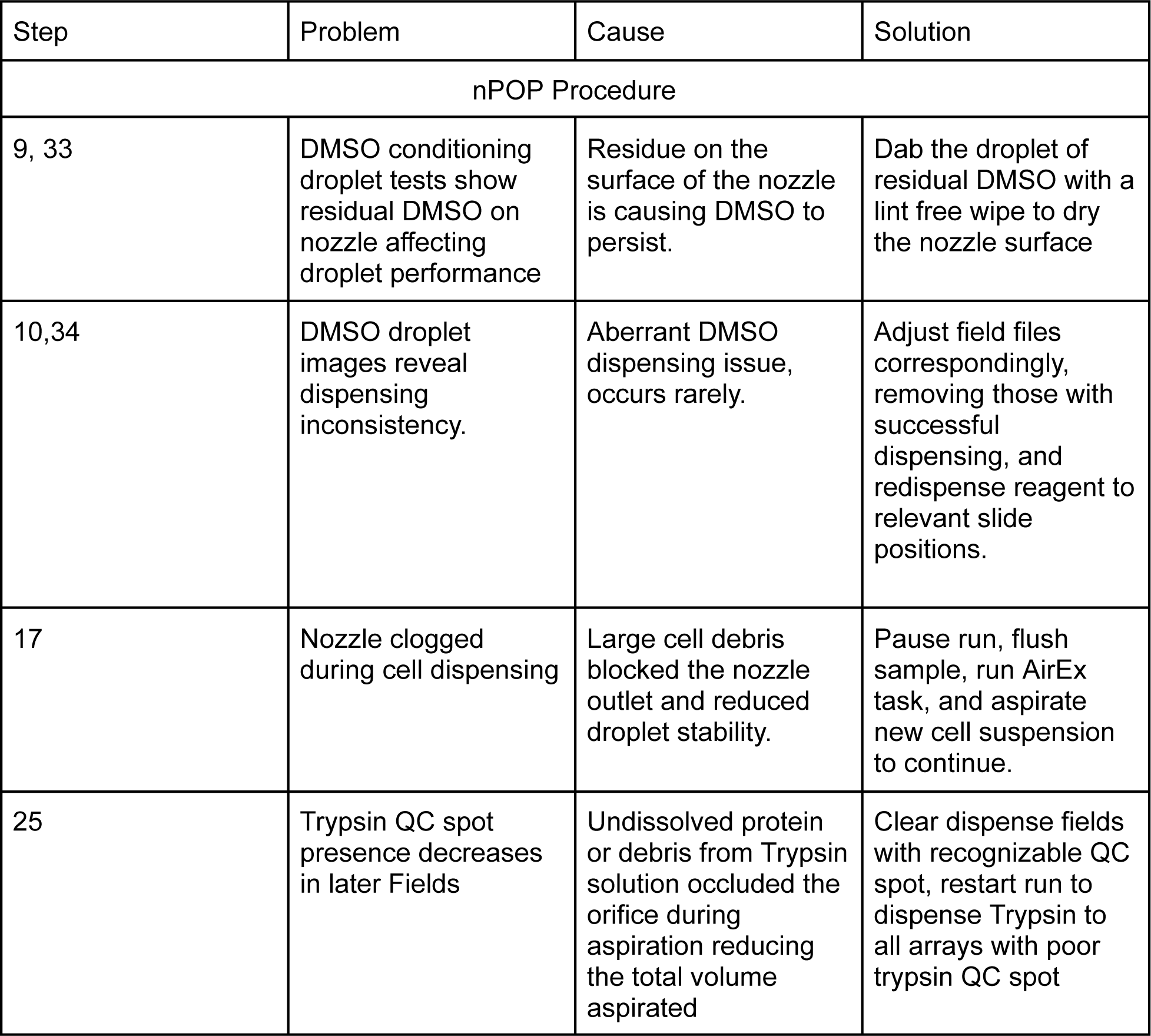
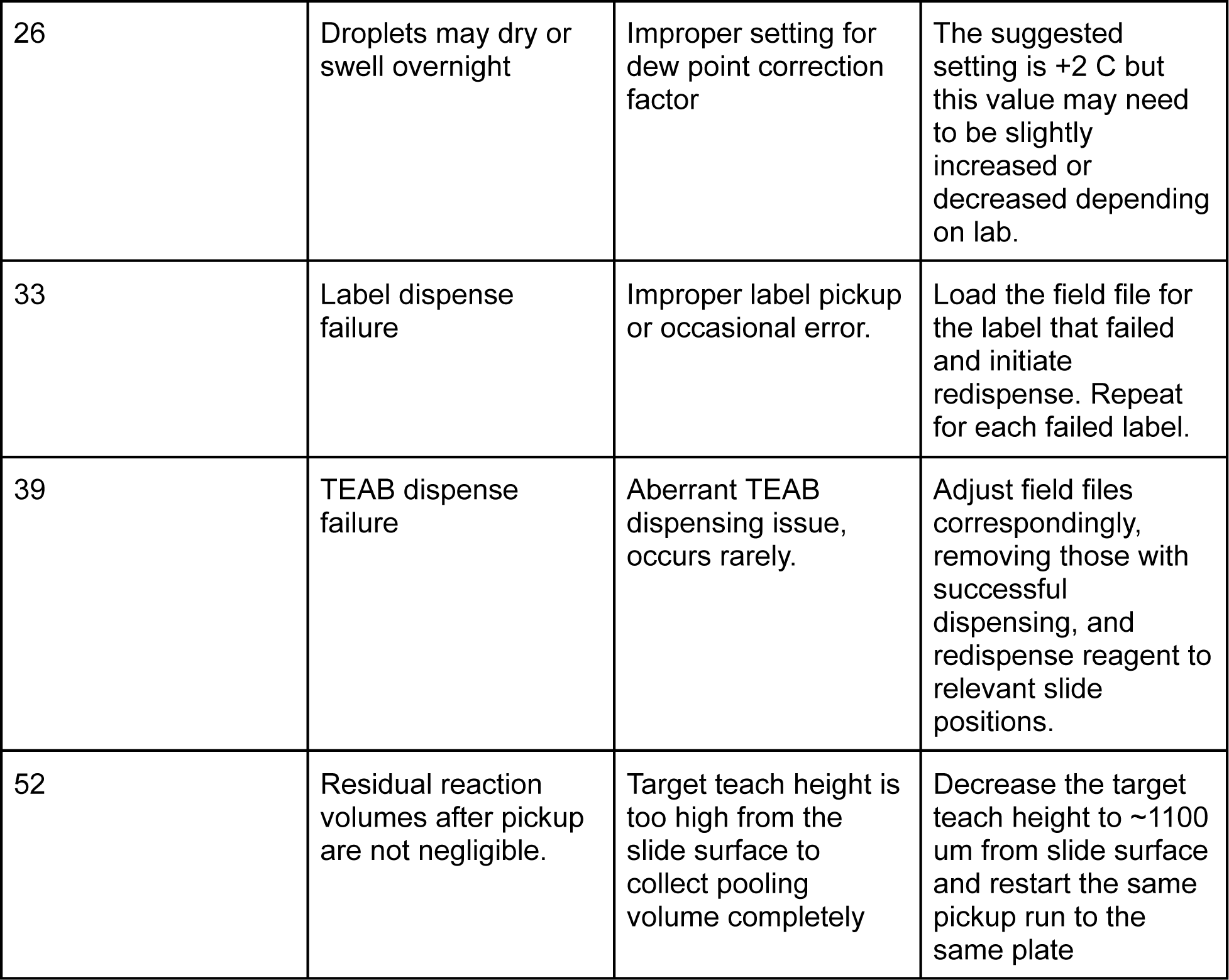
Troubleshooting suggestions for possible problems at each of the indicated steps of the nPOP protocol.

| Step | Problem | Cause | Solution |
| --- | --- | --- | --- |
| nPOP Procedure |  |  |  |
| 9, 33 | DMSO conditioning droplet tests show residual DMSO on nozzle affecting droplet performance | Residue on the surface of the nozzle is causing DMSO to persist. | Dab the droplet of residual DMSO with a lint free wipe to dry the nozzle surface |
| 10,34 | DMSO droplet images reveal dispensing inconsistency. | Aberrant DMSO dispensing issue, occurs rarely. | Adjust field files correspondingly, removing those with successful dispensing, and redispense reagent to relevant slide positions. |
| 17 | Nozzle clogged during cell dispensing | Large cell debris blocked the nozzle outlet and reduced droplet stability. | Pause run, flush sample, run AirEx task, and aspirate new cell suspension to continue. |
| 25 | Trypsin QC spot presence decreases in later Fields | Undissolved protein or debris from Trypsin solution occluded the orifice during aspiration reducing the total volume aspirated | Clear dispense fields with recognizable QC spot, restart run to dispense Trypsin to all arrays with poor trypsin QC spot |
| 26 | Droplets may dry or swell overnight | Improper setting for dew point correction factor | The suggested setting is +2 C but this value may need to be slightly increased or decreased depending on lab. |
| 33 | Label dispense failure | Improper label pickup or occasional error. | Load the field file for the label that failed and initiate redispense. Repeat for each failed label. |
| 39 | TEAB dispense failure | Aberrant TEAB dispensing issue, occurs rarely. | Adjust field files correspondingly, removing those with successful dispensing, and redispense reagent to relevant slide positions. |
| 52 | Residual reaction volumes after pickup are not negligible. | Target teach height is too high from the slide surface to collect pooling volume completely | Decrease the target teach height to ~1100 um from slide surface and restart the same pickup run to the same plate |

## Timing

The nPOP sample preparation can be completed in one full day, but is best split across two experiment days.

### nPOP Day 1: 2-4 hours

● Instrument startup and priming: 30 minutes
● Hands-on time for starting the instrument, sorting cells, and dispensing reagents required for starting the instrument and dispensing cells and reagents is about 1-4 hours depending on the number of cells sorted. Sorting around 3,000 cells will push the time to about 3 hours.

### nPOP Day 2: 2-4 hours

● On day two cells are labeled and transferred to a 384 well plate for LC/MS analysis. Hands on time preparing and sorting the labels and starting the pickup process is about 3 hours total.
● Cleaning nozzles and shut down: 30 minutes

## Box 1: Additional resources

● **Video tutorials** on performing nPOP: https://scp.slavovlab.net/nPOP
● **Community Guidelines** for single-cell MS proteomics^1^: https://single-cell.net/guidelines
● **Protocols** for preparing and optimizing LC-MS/MS platforms for single-cell MS proteomics^1,39,58^: https://scp.slavovlab.net/protocols
● **Computational tools** for single-cell proteomics data: https://scp.slavovlab.net/computational-analysis

## Anticipated results

The nPOP method has been applied to study protein covariation across U937 monocyte and WM-989 cancer cell lines^7^, pancreatic beta cell differentiation^30^, bone marrow derived macrophages^28^ and other systems. All of these studies exemplify anticipated results, and here we provide another example with a smaller data set designed to facilitate introduction to the protocol. To facilitate the reproduction of the protocol example, we used single cells and bulk samples from three accessible cell lines representing different cell types. The bulk samples are included to demonstrate technical benchmarks for the assessment of nPOP^1,6^.

The optimization of MS data acquisition parameters for single-cell analysis can be performed with standards and has been detailed in ref.^39,56,57^. After this optimization, the success of the sample preparation can be assessed with the quality control (QC) reports generated by the QuantQC package in R. The full QuantQC report can be found in supplemental file 1, and several plots are highlighted in Fig. 3.

In multiplexed workflows, it is important to assess the signal strength relative to the background noise, which may originate from contaminants or suboptimal labeling. This can be quantified by comparing the intensities measured from single cells relative to the ones from negative control channels that receive all the same reagents but without a single cell. To provide such an assessment, QuantQC plots the sum of intensities from negative controls and real single cells, Fig. 5a. The results indicate that the intensities corresponding to the single cells is 10 fold higher than from the negative controls, and completely eliminated when peptide identifications are filtered at channel q-values below 1%. Such stricter quality filtering on q values for plexDIA^17^ or MS2 spectral purity for SCoPE2^6^ is recommended when minimizing background influences is more important than optimizing proteome coverage.

**Figure 5.**
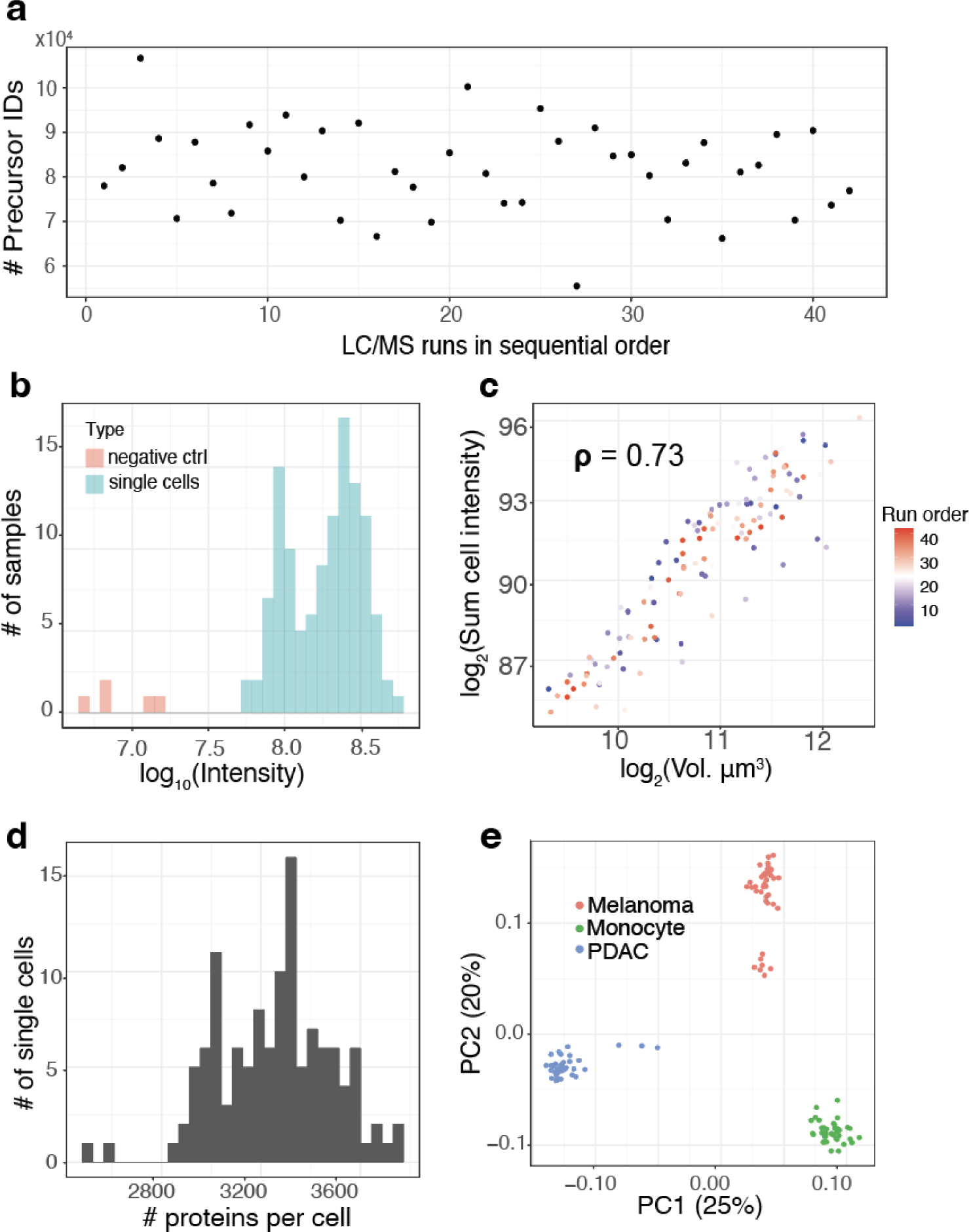
Comparison of throughput and reaction volume. **a**, Number of identified precursors over the course of the LC-MS/MS runs. The numbers remained stable, indicating stable data acquisition. **b**, Distribution of the total signal (estimated as summed intensity from all peptides) for both single cells and negative controls, which have received trypsin and label but no cell. **c**, Cell volume has strong positive correlation with summed peptide signal as a proxy for total protein content indicating consistency of sample preparation. **d**, Distribution of the number of proteins quantified per cell. **e**, Principal component analysis shows that cells discretely cluster by cell type. The 2 clusters of melanoma cells correspond to previously characterized subpopulations in this cell line^7^.

Once it has been established that the single-cell signal has been quantified at a level greater than the background signal, QuantQC plots identifications and ion intensities across LC/MS runs in time to check for systematic trends in LC/MS quality that could induce batch effects in data, Fig. 5b. However, even when performance remains stable over time there can be run to run variance reflecting inconsistent sample preparation recovery or quantity. Thus, consistency can be further measured by plotting the total amount of protein recovered versus the measured cell volume, Fig 5c. This correlation becomes less reliable for cells with smaller sizes (with diameters below 12 microns) because of reduced accuracy of diameter measurements near the lower limit of cellenONE. Nonetheless, the metric usually provides a useful indication for sample preparation and LC/MS consistency if cells are in the range 13-14 microns and up.

QuantQC reports the number of proteins quantified in individual single cells, Fig. 5d, and the number of peptides identified per cell, supplemental file 1. In this data set, an average of 3,202 proteins and 24,347 precursors were identified per single cell. The relationship between the analyzed single cells is visualized by principal component analysis in the space of all proteins identified in at least 10 cells (4,619 proteins) is shown in Fig. 5e. However, separation in low dimensional space does not necessarily reflect accurate measurements, as these trends could arise from batch effects. Thus the QuantQC package color codes cells with experimental factors that may contribute to batch effects and artifacts. These plots are generated automatically as part of a standard html report available as supplemental file 1.

To evaluate the quantitative accuracy of the single cell measurements, we compared single-cell protein fold changes between cell types to the corresponding fold changes estimated from bulk samples analyzed by label-free DIA. The single-cell fold changes were averaged across single cells as performed previously ^6,17^ and suggested by the community recommendations^1^. The fold-change correlations range from 0.77 to 0.82, as shown in Fig. 6a,b,c. These correlations demonstrate that protein quantification from multiplexed single-cell proteomics using nPOP is highly consistent with quantification from conventional non multiplexed methods.

**Figure 6.**
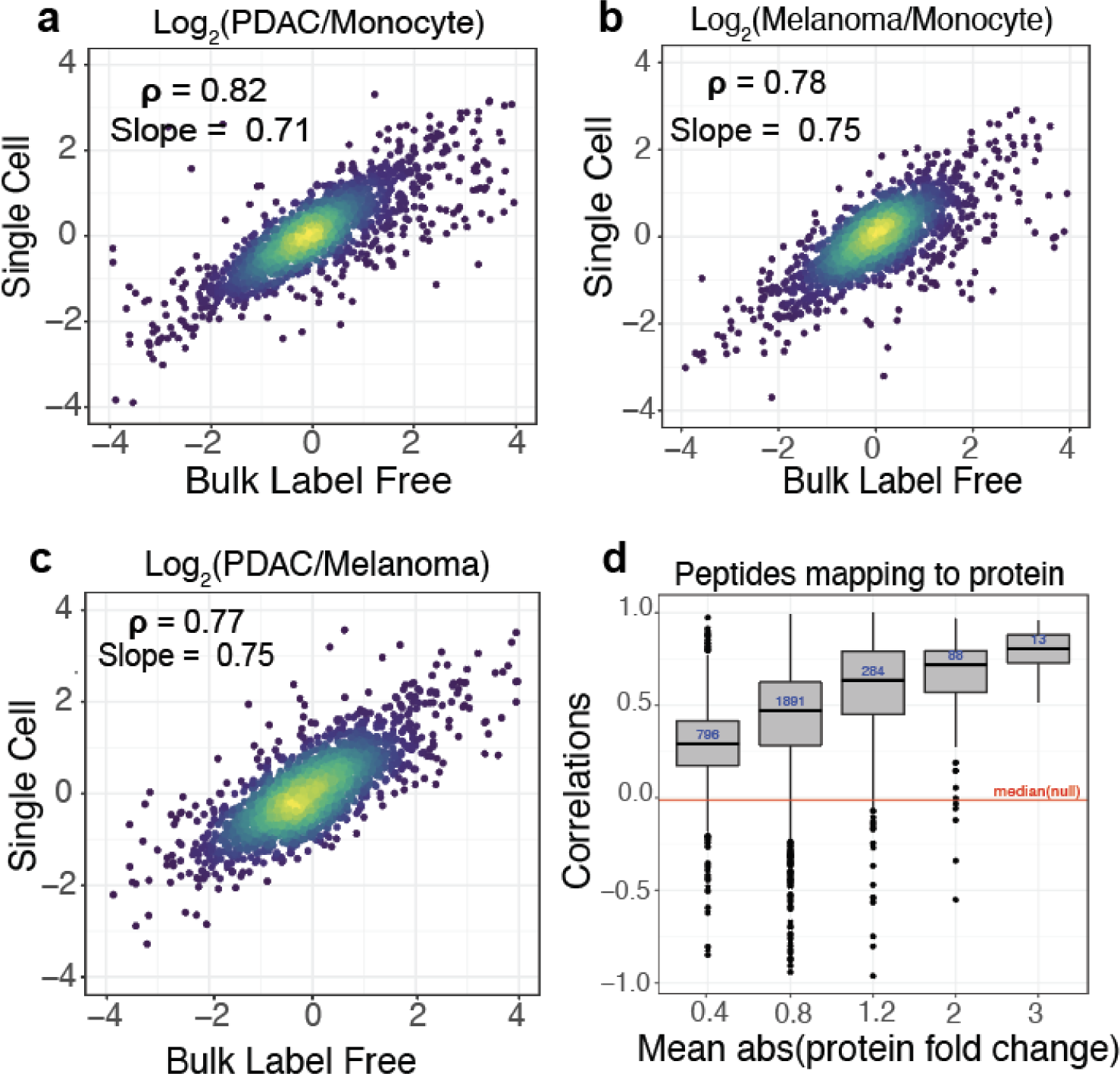
Evaluating quantitative accuracy. All pairwise protein fold changes between the 3 cell types were estimated from single-cell plexDIA measurements using nPOP and from bulk samples analyzed by label free DIA. The corresponding estimates were compared on a log2 scale: **a**, PDAC / Monocyte **b**, Melanoma / Monocyte and **c,** PDAC / Melanoma. For each pair of cell types, single-cell fold changes were averaged *in silico*^1^. **d**, Consistency of protein quantification is estimated by the correlations among peptides mapping to (and thus likely originating from) the same protein. The distributions of these correlations are binned by the absolute fold change variation of the proteins. Proteins varying more across the single cells have higher correlations. The red line represents the median of the null distribution of correlations computed between peptides from different proteins. This plot is generated by QuantQC.

In addition to correlating corresponding fold changes, we sought to quantify the dynamic range of the single-cell measurements. To this end, we computed the slope of the line between fold changes measured in single cells and bulk using total least squares. The slope of the line quantifies the degree to which the magnitude of the observed fold changes are compressed in single cells relative to those measured with bulk methods where quantitative accuracy has been previously validated. The slopes are close to 1, which suggests only a slight ratio compression, Fig. 6a,b,c. However, the dynamic range of old changes, over 100-fold, is preserved in single cells. The slight fold-change compression may arise for various reasons, including interferences, or lowly abundant peptides below the limit of detection in single-cell measurements. The latter challenge may be mitigated by improving the handling of missing data^59^.

Lastly, in the absence of bulk measurements, an internal assessment of the accuracy of MS measurements can be obtained by correlating relative peptide levels across single cells for peptides that map to the same protein. This correlation depends on the signal (biological variance across cells) to noise ratio, and thus its strength depends on the variance of the underlying protein across single cells. QuantQC plots the distribution of correlations faceted by the average absolute fold change of the corresponding protein as a measure of variance, Fig 6d.

## Supplemental Data

Supplementary file 1 is the full QuantQC report. It is available at https://scp.slavovlab.net/nPOP and from <u>Supplementary file 1</u>. All raw and processed data are available at MassIVE MSV000093494 and via the nPOP website: https://scp.slavovlab.net/nPOP.

## Software and Code Availability

The QuantQC package is available at https://github.com/SlavovLab/QuantQC. The most updated nPOP specific software and custom accessories are available through the nPOP Partnership Program offered by Cellenion. Reach out to Joshua Cantlon for sign up information.

## Supporting information

Supplementary file 1, which is the full QuantQC report.

## Acknowledgements

We thank members of the Slavov laboratory for discussions and comments. This work was supported by an Allen Distinguished Investigator award through The Paul G. Allen Frontiers Group, and an R01 award from the NIGMS of the NIH to N.S. (R01GM144967), a MIRA award from the NIGMS of the NIH to N.S. (R35GM148218) and a UH3CA268117 award to N.S.

