## Supplementary file 1, which is the full QuantQC report. for "Massively parallel sample preparation for multiplexed single-cell proteomics using nPOP"

Quant DO-MS


### Quant DO-MS

#### 11/29/2022

### Results

#### LC/MS Performance Statistics

##### IDs and intensities

##### Retention time drift

#### Single Cell Statistics

##### Negative control / single cell comparison

Here we plot the summed intensities across all peptides from a given
single cell, for all single cells and negative controls in the data.

##### Carrier Statistics

```
## [1] "No carrier used"
```

##### Protein and peptide numbers per cell

All the peptides plotted are identified at 1% global FDR.

##### Data completness based on filtering (uses intersected subset of proteins from channel qval filtered data)

##### Correlations between peptides mapping to protein compared to variance

##### Cell size information

##### PCA and batch effects checks

##### Additional dimensionality reduction and clustering

##### Viz proteins of interest

```
## [1] "No Protein added"
```

#### CellenONE\_info

The well position each set is transported into is overlayed in the
middle of the cluster of cells that compose a labeled set.

##### Cell type positions across slide

##### Label position across slide
